# TAXON INCOMPLETENESS AND DISCRETE TIME BINS AFFECT CHARACTER CHANGE RATES IN SIMULATED DATA

**DOI:** 10.1101/2020.07.09.194787

**Authors:** Jorge R. Flores

## Abstract

Estimating how fast or slow morphology evolves through time (phenotypic change rate; PR) has become common in macroevolutionary studies, and has been important for clarifying key evolutionary events. However, the inclusion of incompletely scored taxa (e.g., fossils) could affect PR estimates and potentially mask real PR patterns. This effect being also affected by the usage of arbitrary discrete time bins. Here, the impact of taxon incompleteness (unscored data) on PR estimates is assessed in simulated data. Three different time bin series were likewise evaluated: bins evenly spanning the tree length (i), a shorter middle bin (ii), and a longer middle bin (iii). The results indicate that PR values decrease as taxon incompleteness increases. Statistical significant PR values, and the dispersion among PR values, depended on the time bins. These outcomes imply that taxon incompleteness can undermine our capacity to infer morphology evolutionary dynamics and that these estimates are also influenced by our choice of discrete time bins. More importantly, the present results stress the need for a better approach to deal with taxon incompleteness and arbitrary discrete time bins.

## Introduction

Along with unravelling phylogenetic patterns, another goal of comparative and evolutionary biology is estimating how slow or fast character systems have evolved in different lineages through time [1–5]. Assessing morphology change rate (phenotypic change rate, “PR”) improves our understanding of evolutionary events and helps to explain morphological diversity. PR estimates, for instance, were found to be coupled with morphological disparity and phyletic diversity in certain taxonomic groups [6,7]. Furthermore, in both extant and extinct groups, PRs were shown to be linked with events of adaptive radiations and climatic changes [8,9]. The complex nature of morphological data, however, imposes challenges to infer PR in some organisms [10].

Incompletely scored taxa characterise many morphological datasets, and include two types of unscored entries: missing data (“?”) and inapplicable characters (“-”). Both can be high in matrices that consist of fragmentary or poorly preserved fossils [11] and can reduce taxon stability during phylogenetic inference [e.g., 12]. Some methods to estimate PR treat both unscored data types as “NA” [4] while their treatment in others approaches – designed to infer divergence time based on molecular data – is rather unclear [13]. Yet, some authors have preferred to employ the latter to calculate PR [e.g., 14]. In Claddis [4], an R package, PRs are derived from the number of character state changes occurring along a branch of a certain length. Subsequently, PR values are deemed high or low relative to other data partitions (time bins, branches, etc.) based on a Likelihood Ratio test. Although branch incompleteness is considered in estimates of PR, there is still no satisfactory approach to deal with missing data or inapplicable characters, and their inclusion can lead to false negatives (i.e., not recognising PRs statistically lower or higher than the remaining data partitions [4]).

A further issue with taxon incompleteness (either missing data or inapplicable states) is that it may depend on several factors other than just fossil preservation (e.g., the ability of the observer to recognise and score certain characters). As taxonomic studies advance, a reasonable expectation is that the amount of unscored data change over time and for a given taxonomic group. At present, taxon incompleteness is known to influence PR estimates [15], though its precise impact is still uncertain since no quantitate evaluations were conducted [4,15–17]. Additionally, studies of morphology evolutionary dynamics often involve the assessment of PR in discrete time bins (e.g., [6]). However, accounting for the influence of taxon incompleteness in different time bins is not straightforward given that branches might span through multiple time bins [10]. Thus, the PR for a given time bin depends not only on branch completeness but also on branch lengths and time bins.

In this study, the impact of taxon incompleteness on PR estimates is evaluated based on simulated data. Although simulated data oversimplify many of the main aspects of empirical data, factors possibly affecting PR estimates other than taxon incompleteness (e.g., polymorphism) are easier to control as compared to real datasets. The influence of unequal time bin lengths on PR is also explored. The outcomes shed light on the possible restrictions posed by taxon incompleteness in our understanding of morphological evolution.

## Methodology

Ten initial datasets consisting of 30 taxa and 300 binary characters were simulated in TNT 1.5 [18] under a modified Jukes-Cantor (JC) model wherein character states were equiprobably distributed among matrix cells. Datasets were simulated using random trees as model with branch lengths varying from 0.1 to 100. In the simulated matrices, scored cells were subsequently replaced by missing data (“?”). To do so, first, a maximum percentage between 27% and 50% of the original scored cells was set for being replaced with missing data in the initial datasets. For each initial matrix, 10 iterations were then performed wherein an increasing number of the missing cells were solved (choosing randomly between “0” and “1”) until less than 1% of missing data remained in the matrices. This generated 10-matrix series, simulated upon the same model tree, wherein the only factor that varied was the amount of unscored data. This procedure was applied in the 10 initial simulated matrices; thus, concluding in 100 matrices for subsequent estimation of PR. Although larger matrices (e.g., > 500 characters) would increase the power of PR estimations, smaller datasets (e.g., < 100 characters) reduce the computational effort. Mid-sized matrices, like those analysed here, represented a compromise between both efficient computational effort and statistical power. An extended explanation of these procedures, along with the TNT scripts, are provided as supplementary material.

Phenotypic change rate (PR) was assessed in the simulated matrices by using three time bins and using the function “DiscreteCharacterRate” of the R package Claddis [4,19]. PRs were estimated upon the model trees and calculated as the average number of character changes per branch length and branch completeness [4]. Studies involving PR commonly use discrete time bins of different length. To evaluate the effect of using different bin lengths, three series of time bins were evaluated (*i-iii*). In these series, the three time bins spanned different proportions of the total tree length. Firstly, time bins were each set to span 0.3 of the entire tree length (i.e., time bins of equal length; *i*). Secondly, a shorter middle bin was defined by spanning from 0.3 to 0.25 of the tree length (i. e, a five per cent; *ii*). Thirdly, a longer middle bin was delimited extending from 0.85 to 0.15 of the tree length (i.e., 70 per cent, *iii*). To take into account branch incompleteness per time bin, the “Lloyd” option from the “DiscreteCharacterRate” function was employed (see details in Llyod [4] and Claddis documentation). Finally, both PR values per bin and the proportion of statistically high or low PRs were plotted against the proportion of missing data for the three time bin series considered (*i-iii*).

## Results

The analyses showed that, across the 100 simulated datasets, PR values diminished as the proportion missing data increased (figure 1). For these JC simulated datasets, PR values in the first time bin tended to be statistically higher than the remaining bins (red dots, figure 1). Conversely, in the remaining bins, PR values were statistically lower than in the other data partitions (blue triangles and squares, figure 1). Comparatively, the initial fully-scored datasets rendered a similar pattern among bins, though with higher proportion of statistically significant PR values (figures S1, Supplementary Material). Thus, adding unscored data enhanced the effect caused by different bin lengths.

**Figure 1.**
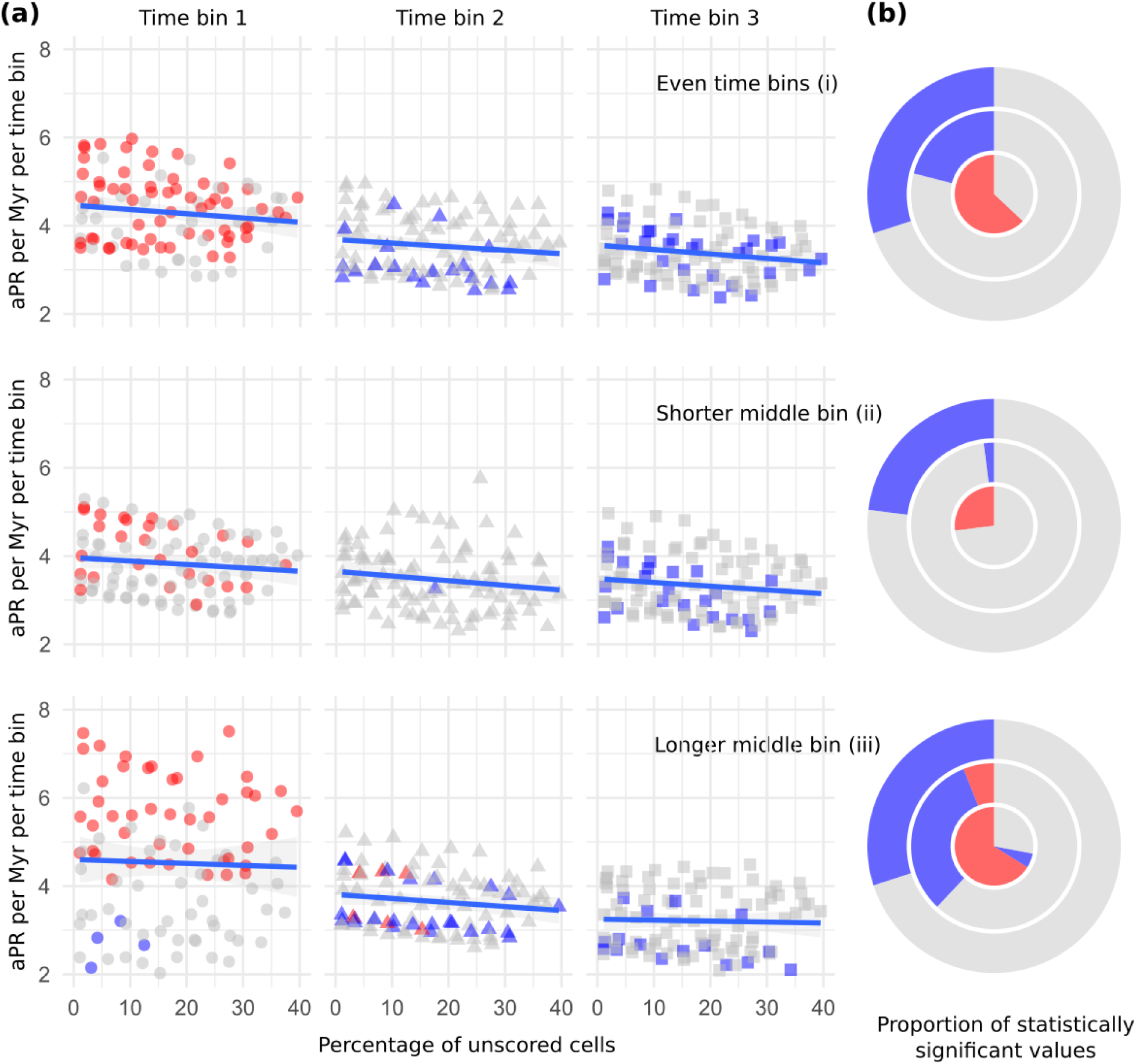
Average number of character state changes (average phenotypic change rate, aPR) per Myr in each time bin against the percentage of unscored cells (*a*). The proportion of statistically low or high PR values per bin, across 100 simulated datasets, is also shown as bullseye plots wherein time bins are depicted as rings (the innermost being the first time bin; *b*). Red and blue symbols indicate statistically high and low values, respectively. Grey represents no statistical significance.

Varying the extension of time bins affected the PR estimates (*i-iii*, figure 1*b*). As compared to even time bins (*i*), uneven time bin series (*ii-iii*) altered observed PR values, the dispersion among PR values within time bins and the proportion of statistically high or low PR values (figure 1). When the middle time bin was shorter (*ii*), the dispersion among PR values raised (from 2.1 to 5.8, figure 1*a*), and the proportion of significant PR values was minimised as compared to the remaining time bin series (figure 1*b*). Upon extending the length of the middle bin (*iii*), PR response to the percentage of unscored cells was slightly faster than in the other middle bins (slope of the tendency line, figure 1*a*). Likewise, the dispersion among PR values was minimised (figure 1*a*) and the proportion of significant values was maximised (either low or high, figure 1*b*). Additionally, it must be noted that PR values within the first time bin were also affected after modifying the extension of the second time bin. Extending the middle time bin (i.e., shortening the first time bin) increased the dispersion among PR estimates (*iii*, figure 1*a*) while the proportion of statistically significant values was reduced when the first time bin was longer (*ii*, figure 1*b*). PR values within the third time bin were stable to the variation of time bin lengths (*i-iii*, figure 1).

## Discussion and Final remarks

The present estimations of the number of character change relative to branch lengths (phenotypic change rate, PR) were sensitive to both taxon incompleteness and the selected time bins (figure 1). Taxon incompleteness has been long studied in the context of phylogenetic inference (e.g., [20]), though its impact on PR estimation is not fully certain. While some authors have excluded characters or taxa that were highly incomplete [21,22], taxon incompleteness has not been explicitly addressed in other studies (e.g., [7,14]). In early tetrapods, the inclusion of fragmentary fossils was argued to exaggerate both fast and slow PRs [17]. Following this logic, PR should be expected to increase their statistically high or low values as more unscored data is sampled. However, in the present analyses, the response of PR values and the proportion of unscored data was invariably negative (figure 1*a*). Moreover, if incompleteness would accentuate PR values [17], estimates in the first time bin should have increased as more unscored cells are present. Instead, the opposite pattern is seen regardless of the PR statistical significance (figure 1*a*). The decay in PR observed here as a consequence of unscored data was also seen for disparity-through-time (DTT) metrics [23]. Overall, this result expands upon previous analyses that showed the negative impact of incompleteness on disparity measures [23], and indicate that the inclusion of highly fragmentary taxa could hinder the distinction of an otherwise high phenotypic rate.

Another interesting outcome is the effect of varying the limits of the middle time bin on the statistical significance of the PR values (figure 1*b*). As already mentioned, addressing branch incompleteness is not straightforward given that branches commonly span time bin limits (see [4] and Claddis R package documentation). In DTT studies, it has been already noticed that the usage of discrete time bins can introduce biases [23,24]. Time bins are commonly based on stratigraphic data and, therefore, are of unequal lengths (e.g., [6,17,25]). In those cases, biases are due to the uneven number of taxa included in different time bins: disparity is higher in longer time bins wherein more taxa are included [23,24]. Although rarely discussed, bin lengths can also affect the statistical power of the PR test. This is because more branches are to be sampled in longer time bins. Therefore, as the sampling size increases, the statistical power of the test improves. Conversely, when time bins are shorter, fewer branches are sampled and the statistical power is reduced. In the present analyses, the proportion of significant low rates in the middle time bin was reduced after shortening the bin limits as compared to longer middle time bins (figure 1*b*). However, it is worth noting that the number of branches present in a specific bin also depends on the topology of the tree: symmetric trees are likely to have more branches included in a time bin as compared to asymmetric trees. Such factor could account for the higher proportion of statistically high PRs in the shorter first time bin (*iii*, figure 1*a*).

In actual matrices, taxon incompleteness involve both missing data and inapplicable character states. Although both data types are different in nature, these are commonly treated equally in current implementations to evaluate PR (e.g., [4]). To take into account incompleteness, both weighting (“Close”, [4, 10]) and “subtree” (“Lloyd”, see details in [4]) approaches were proposed. Another proposal involves down-weighting characters that “concentrate” a large number of the observed changes [4], in a similar way to implied weighting [26]. Nevertheless, since missing data (“?”) and inapplicable characters (“-”) are treated equally, these approaches can overestimate incompleteness. A potential way to discriminate both data types is based on the usage of step-matrix characters. If considering inapplicable character states as additional states, prohibitive transformation costs between these and scored character states could be employed to avoid treating inapplicable data as missing (“NA”). Such an approach requires no *ad hoc* measures (e.g., weighted means, [10]) since inapplicable character state changes are inferred only between terminal nodes and their last ancestor node. Branch incompleteness is thus due to missing data (“?”) or ambiguous character state reconstruction. However, because step-matrix characters are not yet supported in Claddis, implementing this strategy is not possible.

To conclude, the present study shows that taxon incompleteness (unscored data) reduces PR estimates. Likewise, the choice of different time bin lengths modifies our capacity to observe statistically significant values. These outcomes stress the need for approaches that could deal with unscored data and avoid biases emerging from the definition of discrete time bins. Simulated data here was conceived to render statistically significant values in all the time bins. Although simulated data do not take into account the complexity of real data, it allows controlling other factors. Further studies should focus on the impact of unscored data and discrete time bins in empirical datasets.

## Data accessibility

Scripts for simulating data in TNT and PR estimation in R are available as online supplementary material.

## Acknowledgements

TNT is freely available to users thanks to the Willi Hennig Society.

